# A proteomic study of the dual oncogenic and tumor-suppressive roles of SIRT3 in lung and breast cancer cell lines

**DOI:** 10.1101/2025.08.26.672455

**Authors:** Marisol Ayala Reyes, Diana Lashidua Fernández Coto, Ramiro Alonso Bastida, György Marko-Varga, Jeovanis Gil, Sergio Encarnación Guevara

## Abstract

Mitochondria play a crucial role in metabolism and energy production by generating adenosine triphosphate (ATP) through oxidative phosphorylation. They also help maintain intracellular calcium levels, facilitate communication between the nucleus and cytoplasm, detoxify reactive oxygen species (ROS), and regulate apoptosis (programmed cell death). Reversible acetylation of mitochondrial proteins is a key post-translational modification influencing these processes, with the NAD^+^-dependent deacetylase SIRT3 being a major regulator. While SIRT3 has been described as a tumor suppressor in some contexts and as a tumor promoter in others, its role appears to be tissue- and metabolism-specific. Here, we compared the proteomic and acetylomic responses of lung adenocarcinoma (A549) and breast adenocarcinoma (MCF7) cell lines to SIRT3 inhibition by 3-(1H-1,2,3-triazol-4-yl)pyridine (3-TYP). The two lines were selected based on distinct metabolic phenotypes and reported differences in basal SIRT3 abundance. Total proteome and mitochondrial-enriched fractions were analyzed separately for each cell line to avoid cross-line normalization bias. We identified 6,457 proteins and 4,199 acetylated peptides, revealing divergent pathway enrichments and acetylation patterns following SIRT3 inhibition. A549 cells showed enrichment of oxidative metabolism pathways, whereas MCF7 cells exhibited features consistent with metabolic reprogramming. These findings suggest that the cellular context determines the proteomic consequences of SIRT3 modulation and highlight the importance of metabolism in shaping SIRT3-associated phenotypes.

All raw mass spectrometry data are publicly available in [repository accession number].

**Significance:** SIRT3’s impact on cancer metabolism is context-dependent. By separately profiling lung (A549) and breast (MCF7) cancer cells, we demonstrate that SIRT3 inhibition leads to distinct proteomic and acetylomic outcomes in each cell type. Our findings underscore the need for cell-type-specific strategies when targeting SIRT3 in cancer therapy and provide an open-access proteomic resource for further mechanistic studies.

## 1. Introduction

Mitochondria are essential for cellular energy homeostasis, generating ATP via oxidative phosphorylation (OXPHOS) and coordinating calcium signaling, reactive oxygen species (ROS) balance, and apoptosis. Alterations in mitochondrial function are linked to cancer, cardiovascular disease, and neurodegeneration [1]. Among the mechanisms that regulate mitochondrial activity, reversible acetylation of lysine residues in mitochondrial proteins is particularly important. This post-translational modification can be catalyzed by lysine acetyltransferases (KATs) [2,3], yet no dedicated mitochondrial KAT has been identified to date. Instead, evidence suggests that a substantial fraction of mitochondrial acetylation occurs non-enzymatically, driven by the high concentration of acetyl-CoA (ranging from 0.1 to 1.5 mM) and the alkaline pH (7.9–8.0) of the mitochondrial matrix [4]. Acetylation generally inhibits mitochondrial enzymes, whereas deacetylation by Zn^2+^-dependent deacetylases or NAD^+^-dependent sirtuins can restore activity. Within the mitochondria, deacetylation is primarily facilitated by three sirtuins: SIRT3, SIRT4, and SIRT5. Among these, SIRT3 exhibits the most significant deacetylase activity [5]. SIRT3 targets various proteins and enzymes by removing the acetyl group, which are essential for the proper functioning of mitochondria within the cell. Examples of these targeted proteins include long-chain acyl-CoA dehydrogenase (LCAD) [6,7], pyruvate dehydrogenase (PDH) [8], acetyl-CoA synthetase 2 (AceCS2), isocitrate dehydrogenase 2 (IDH2) [9], superoxide dismutase 2 (SOD2) [10], and complexes I, III, and V (ATP synthase) of the electron transport chain, among others that are crucial for metabolism [11]. Therefore, it has been observed that mitochondrial protein acetylation is thought to exert an inhibitory function [12], suggesting that deacetylation is vital for maintaining mitochondrial function, which is essential in preventing diseases like cancer.

In cancer, the function of SIRT3 is strongly influenced by context. In some tumors, reduced SIRT3 expression drives metabolic reprogramming toward glycolysis, which supports rapid cell proliferation. Restoring SIRT3 activity in these cases can exert a tumor-suppressive effect. In other cancers, high SIRT3 expression maintains oxidative metabolism and strengthens antioxidant defenses, conditions that can promote tumor growth. This dual behavior has been observed in several cancer types and appears to depend on the basal metabolic profile of the tumor, the characteristics of its microenvironment, and its tissue of origin [13].

Lung cancer and breast cancer illustrate this contrast. Both are the leading causes of cancer-related death in men and women, respectively [14]. In lung cancer, SIRT3 is frequently overexpressed and associated with tumor progression [15,16]. In breast cancer, it is often downregulated, and this reduction correlates with more aggressive disease and poorer prognosis [17–19]. These patterns are consistent with the metabolic profiles of their representative cell lines: A549 lung adenocarcinoma cells exhibit a predominantly oxidative phenotype, whereas MCF7 luminal A breast cancer cells show a greater reliance on glycolysis.

From a therapeutic perspective, these observations suggest that SIRT3 inhibition could be beneficial in lung cancer, whereas activation or restoration of SIRT3 activity may be advantageous in breast cancer. However, there are few direct comparative studies examining how SIRT3 modulation affects the proteome and acetylome in these two tumor types.

In the present work, we examine the effects of pharmacological SIRT3 inhibition in A549 and MCF7 cells using 3-(1H-1,2,3-triazol-4-yl)pyridine (3-TYP), a small molecule with higher affinity for SIRT3 than for SIRT1 or SIRT2, although complete specificity is not achieved. We analyzed total proteomes and mitochondrial-enriched fractions in parallel and assessed acetylation stoichiometry. To avoid biases from direct comparisons between biologically distinct cell lines, each line was analyzed separately, comparing untreated controls with 3-TYP–treated samples.

## 2. Material and Methods

### Biological Material

Lung cancer cell lines (A549) and breast cancer cell lines (MCF7) were used in this study. The cell lines were cultured in RPMI 1640 medium supplemented with 10% fetal bovine serum (FBS) and 1X antibiotic-antimycotic. The cells were maintained at 37°C in a 5% CO2 atmosphere. To inhibit SIRT3, the cells were treated with a concentration of 25 nM 3-TYP for 24 hours prior to cellular protein extraction or mitochondrial enrichment. Control cells were maintained in the same conditions without any treatment.

### Determination of Reactive Oxygen Species

The levels of reactive oxygen species were measured in 12-well culture plates containing MCF7 and A549 cells. Both treated and untreated cells were used when they reached 90% confluence. The cells were exposed to a 1.5 µM solution of MitoSOX Red Mitochondrial Superoxide Indicator and incubated for 30 minutes at 37°C in an atmosphere containing 5% CO2, while kept in the dark. After the incubation period, the cell monolayers were detached and centrifuged at 4,500 rpm for 5 minutes. The resulting cell pellet was washed three times with Hanks Balanced Salt Solution (HBSS), preheated to 37°C. The cells were then counted and adjusted to a concentration of 500,000 cells per sample. As a positive control, some wells were treated with Antimycin A at a concentration of 100 µM for 30 minutes. The samples were analyzed using a BD FACSCanto II flow cytometer (BD Biosciences) equipped with BD FACSDiva software, and the data were processed using FlowJo™ software (version 10.10). Graphs were created using GraphPad Prism 8 and RStudio (version 4.3.1).

### Protein extraction

To extract proteins, six million cells from each cell line were homogenized using a protein extraction buffer composed of 4% SDS, 50 mM DTT, and 100 mM Tris-HCl (pH 8.6). The samples were sonicated for twenty cycles while keeping them on ice. Following this, the extracts were centrifuged at 20,000 x g for 20 minutes, and the supernatant was collected. The total protein concentration was then measured using the Pierce™ 660 nm protein assay kit, following the manufacturer’s instructions.

### SDS-PAGE (Sodium Dodecyl Sulfate Polyacrylamide Gel Electrophoresis) and immunoblotting

Protein extracts were subjected to electrophoresis and then transferred to a polyvinylidene fluoride (PVDF) membrane. After a blocking period of two hours, the membrane was incubated overnight at 4°C with a primary anti-SIRT-3 antibody (Cell Signaling Technology) diluted to 1:1000. Following several wash steps, the membrane was incubated for one hour at room temperature with a secondary antibody conjugated to horseradish peroxidase (HRP), which targets the light chain of rabbit immunoglobulin G (IgG) (Santa Cruz Biotechnology), at a dilution of 1:10,000 under blocking conditions. For protein detection, the maximum sensitivity chemiluminescence substrate (SuperSignal™ West Femto, Thermo Scientific™) was used. The images were scanned using a C-DiGit® Blot Scanner (LI-COR), and densitometry analysis was performed using Image Studio 5.2 software.

### Enrichment of Mitochondria

Mitochondrial enrichment was performed using three 175 cm^2^ cell culture dishes, each reaching 80% confluence under the specified conditions. To separate the cell monolayer, trypsinization was conducted, followed by centrifugation at 208 xg for 5 minutes. After removing the supernatant, the resulting cell pellet was homogenized in 15 mL of HEPES buffer, which contained 20 mM HEPES, 2 mM EGTA, 250 mM sucrose, and 0.5% albumin at pH 7.4). The mixture was then centrifuged at 1,500 x g for 20 minutes at 4°C, and the supernatant was collected. This process was repeated three times, combining the supernatants obtained at each stage.

The combined supernatants were centrifuged at 15,600 x g for 50 minutes at 4°C. The resulting pellet was resuspended in mannitol buffer (210 mM mannitol, 70 mM sucrose, 5 mM EDTA, and 5 mM Tris-HCl) and placed on a sucrose gradient. This gradient was centrifuged in a swing-out rotor (SW41) at 40,000 x g for 60 minutes at 4°C, after which the cloudy phases corresponding to the mitochondrial fraction were collected.

The mitochondrial fraction was then dissolved in mannitol buffer and centrifuged at 18,000 x g for 40 minutes at 4°C. The resulting pellet was mixed with a protein extraction buffer containing 4% SDS, 50 mM DTT, and 100 mM Tris (pH 8.6), sonicated, and quantified before being stored at -20°C for further use. This mitochondrial enrichment process was repeated three times to ensure the reproducibility of the results.

### Chemical Acetylation with NAS-d3

For the chemical acetylation process, we used deuterated N-acetoxysuccinimide reagent (NAS-d3), which was previously synthesized in our laboratory. We began by reducing the disulfide bridges in protein extracts through a 5-minute incubation at 95°C. Next, we alkylated the proteins by adding IAA and allowing the reaction to proceed for 20 minutes in the dark at room temperature. The proteins were then precipitated overnight by adding nine volumes of cold ethanol. After precipitation, we dissolved the proteins in a TEAB buffer containing 1% SDS and 0.5% SDC. They subsequently underwent chemical acetylation in a reaction buffer with NAS-d3, repeating this step twice to ensure a complete reaction. Following acetylation, the samples were deesterified by adding 5% hydroxylamine, followed by another round of ethanol precipitation. Finally, the proteins were digested with trypsin. We accomplished detergent removal by utilizing a mixture of ethyl acetate and TFA, and then thoroughly dried the samples using a speed vacuum.

### LC-MS/MS Analysis and Acetylation Stoichiometry Analysis

The LC-MS/MS analysis was conducted at the Biomedical Center at Lund University in Sweden. The peptides were resuspended under the initial chromatographic conditions and separated using a Dionex Ultimate 3000 RSLC nano UPLC system. This system was online connected to a Thermo Fisher Scientific Q-Exactive HF-X high-resolution mass spectrometer. For the identification and relative quantification of peptides and proteins from the data generated by the LC-MS/MS measurements, MaxQuant software was used.

The data obtained from the LC-MS/MS analysis were utilized to determine the acetylation stoichiometry of identified peptides containing lysine residues. To perform this analysis, we employed the Pview program, which calculates acetylation stoichiometry based on the area of the isotopic distribution in the mass spectrum of the identified peptides.

### Statistical Analysis and Graph Construction in R

To conduct Principal Component Analysis (PCA), we first identified the variability in our data that was due to the “batch effect.” To address this variability, we implemented a normalization procedure to enable accurate clustering. Next, we applied the T-statistic test (Benjamini-Hochberg method) to generate Volcano Plot scatter diagrams. We classified proteins with acetylation levels equal to or greater than 0.4% as over-acetylated proteins in our samples. The significant protein lists from each comparison, along with the lists of over-acetylated proteins, underwent pathway enrichment analysis using GeneCodis [20]. This analysis allowed us to create diagrams illustrating the associations with relevant biological pathways. Finally, we used GraphPad Prism 8 and RStudio (version 4.3.1) to generate graphs.

## 3. Results

### 3.1 Proteomic analysis of SIRT3 in lung and breast cancer cell lines

Our study focused on the independent proteomic analysis of lung adenocarcinoma (A549) and breast adenocarcinoma (MCF7). To assess the function of SIRT3, we utilized the selective inhibitor 3-TYP (3-(1H-1,2,3-triazol-4-yl) pyridine). 3-TYP was used at 25 nM, a concentration near its reported IC_50_ for SIRT3 (16 nM) and well below the IC_50_ values for SIRT1 (88 nM) and SIRT2 (92 nM), indicating higher potency toward SIRT3 under these conditions [21].

Proteins from total cell lysates and mitochondrial-enriched fractions were extracted in triplicate and analyzed by LC-MS/MS (workflow in Figure 1A). By applying a filtering criterion of 70% valid values across all samples, we identified a total of 6,457 proteins. Among these, we detected 4,199 acetylated peptides. The pattern of acetylation stoichiometry is illustrated in **Figure 1B**, where the percentage of acetylation is represented on a scale from 0 to 100%.

**Figure 1.**
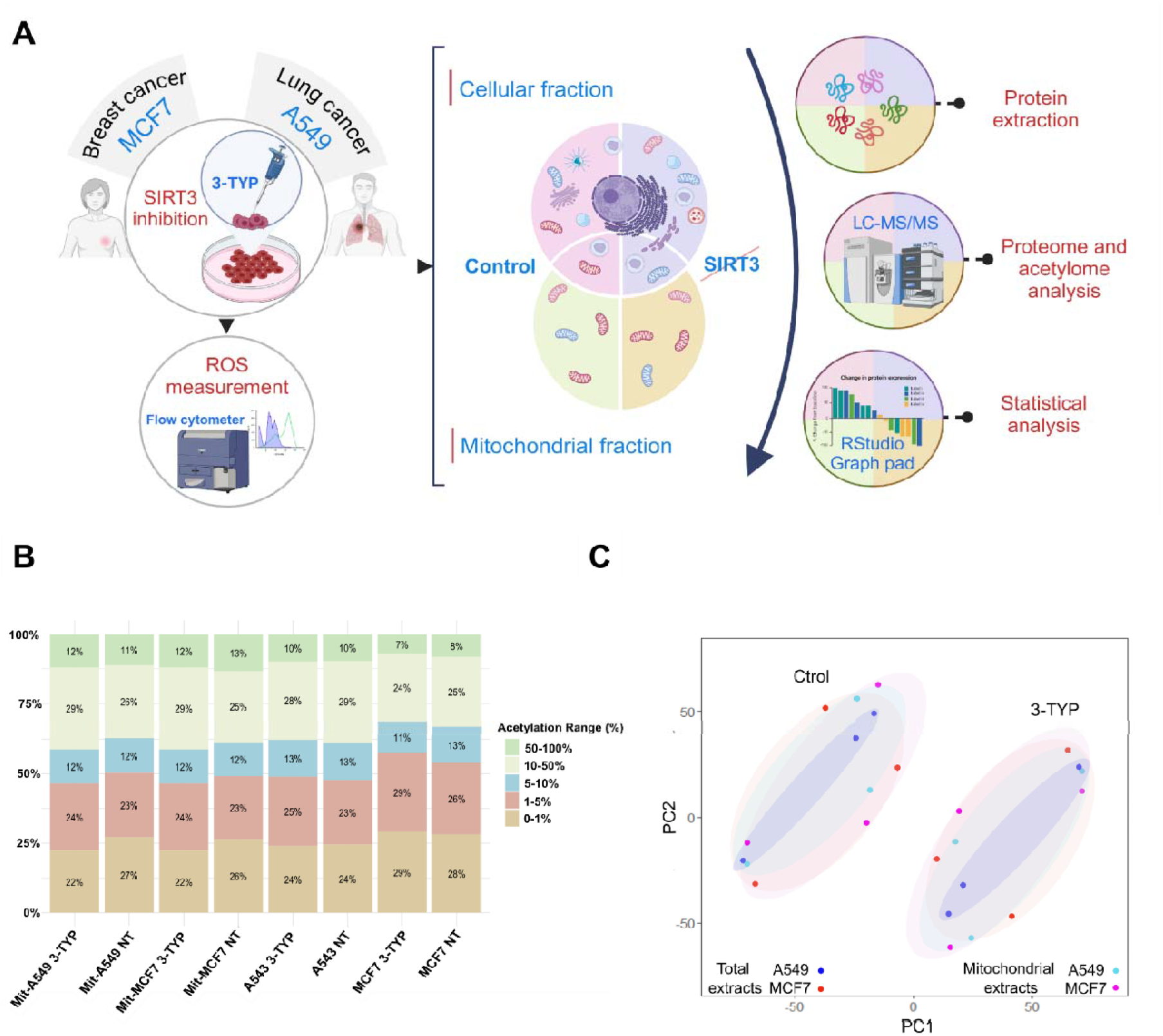
Overview of the workflow and validation for sample grouping using Proteome Principal Component Analysis (PCA). (A) Schematic overview of the experimental design, including cell culture, 3-TYP treatment, protein extraction (total and mitochondrial fractions), and LC–MS/MS analysis. (B) The percentage of acetylation observed in each sample is displayed on a scale from 0 to 100%, categorized into the following ranges: 0–1%, 1–5%, 5–10%, 10–50%, and 50–100%. (C) PCA results for total and mitochondrial samples from A549 and MCF7 cells, highlighting control samples to those treated with the 3-TYP inhibitor.

Principal Component Analysis (PCA) of the cellular and mitochondrial proteomes, after correcting for batch effects, showed clustering of samples based on their response to SIRT3 inhibition, as illustrated in **Figures 1C**.

### 3.2 SIRT3 exhibits contrasting roles in tumor progression between lung and breast cancer

Basal SIRT3 protein levels were quantified independently in each cell line. Triplicate immunoblotting showed that SIRT3 abundance was significantly higher in A549 cells than in MCF cells when normalized to total protein load (Figure 2A). This finding was corroborated by proteomic quantification, which also confirmed higher SIRT3 levels in A549 across both total lysates and mitochondrial fractions (Figures 2B and 2C). These differences are consistent with previous reports describing elevated SIRT3 expression in lung cancer and reduced levels in breast cancer [15,22]. While such variation may relate to the distinct biological roles proposed for SIRT3, it is important to note that these measurements reflect inherent properties of the cell lines and do not, on their own, establish functional roles without further experimental validation.

**Figure 2.**
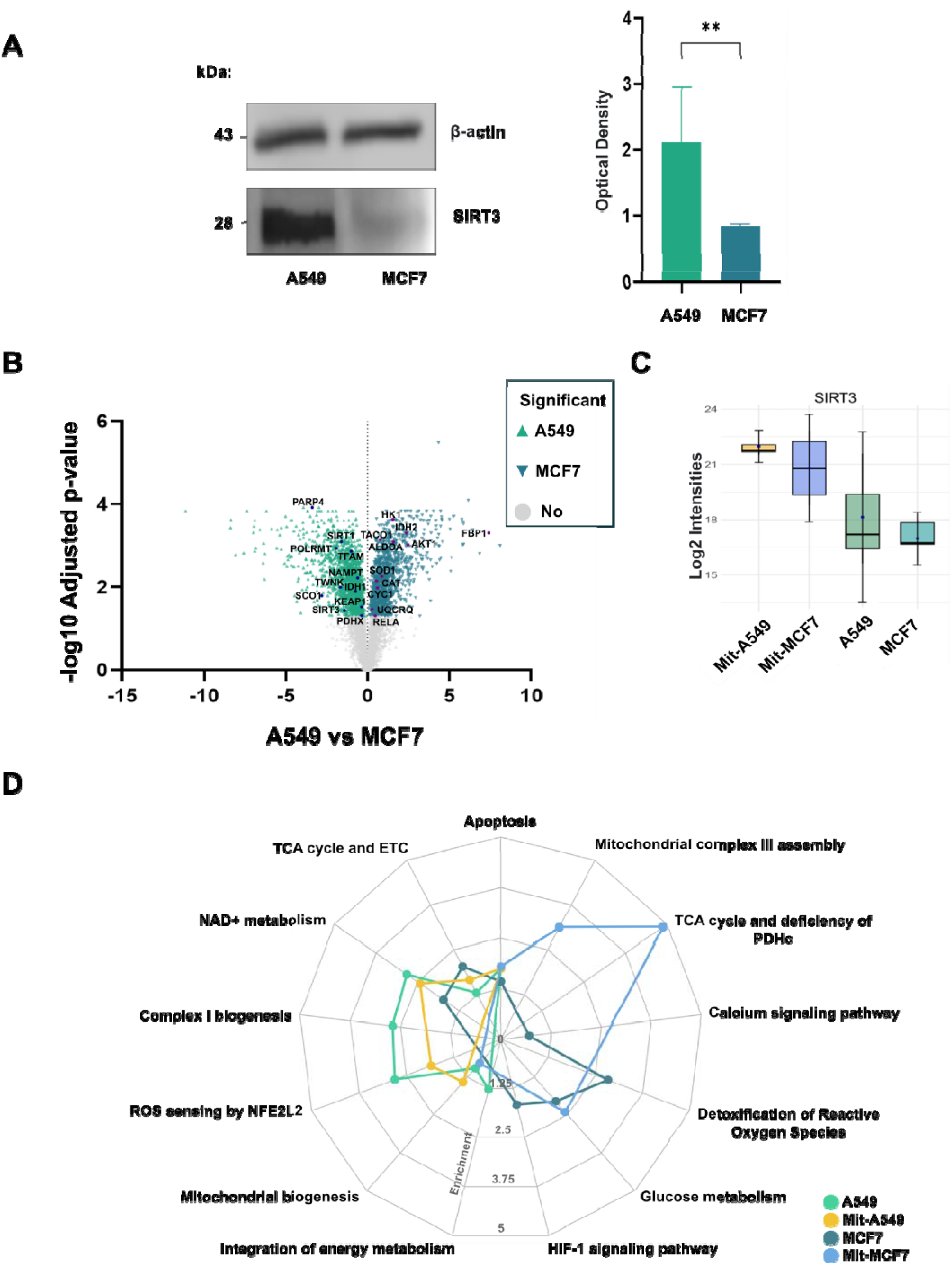
SIRT3 shows different expression levels in the MCF7 and A549 cell lines, with higher expression observed in the A549 cells and lower expression in the MCF7 cells. (A) Representative immunoblot images illustrate SIRT3 expression in both cell lines, along with optical densitometry results derived from triplicate immunoblots for SIRT3. An unpaired t-test was used for statistical analysis, with asterisks indicating significance at P < 0.05. (B) The volcano plot illustrates protein expression between the A549 and MCF7 cell lines, highlighting key proteins. (C) SIRT3 signal intensities were measured using mass spectrometry. (D) Pathways enrichment was performed comparing the proteomes of A59 and MCF7 cells.

Pathway enrichment analyses were performed separately for each cell line to avoid bias from cross-line normalization. In A549 control samples, oxidative metabolism pathways—including mitochondrial biogenesis and nicotinamide adenine dinucleotide (NAD) metabolism—were significantly enriched (Figure 2D). Given that NAD is an essential cofactor for sirtuin activity, these findings align with the higher abundance of SIRT3 and SIRT1 observed in this cell line. In mitochondrial fractions from A549 cells, SIRT3 remained among the most abundant proteins, suggesting that a substantial proportion of its activity may be localized within mitochondria.

In contrast, MCF7 control samples showed enrichment of glycolytic metabolism pathways and HIF-1 signaling, consistent with a more glycolytic phenotype. Mitochondrial fractions from MCF7 cells were enriched for tricarboxylic acid (TCA) cycle components but exhibited reduced representation of pyruvate dehydrogenase complex (PDHc) subunits. As PDHc catalyzes the conversion of pyruvate into acetyl-CoA for entry into the TCA cycle, its lower abundance may indicate altered mitochondrial utilization of glycolytic substrates.

These baseline differences in SIRT3 abundance and associated metabolic pathways provide the context for interpreting the cell-type–specific proteomic and acetylomic responses to SIRT3 inhibition described in the following sections.

### 3.3 Inhibiting SIRT3 in A549 cells may enhance the key pathways that drive disease progression

After confirming the overexpression of SIRT3 in the A549 cell line, we analyzed and compared total and mitochondrial proteins between control samples and those treated with 3-TYP. In the total protein samples from the control group, we found significant enrichment in pathways related to oxidative metabolism, including the TCA cycle, the ETC, the assembly of complex I within the ETC, and transcriptional activation of mitochondrial biogenesis (**Figure 3A**).

**Figure 3.**
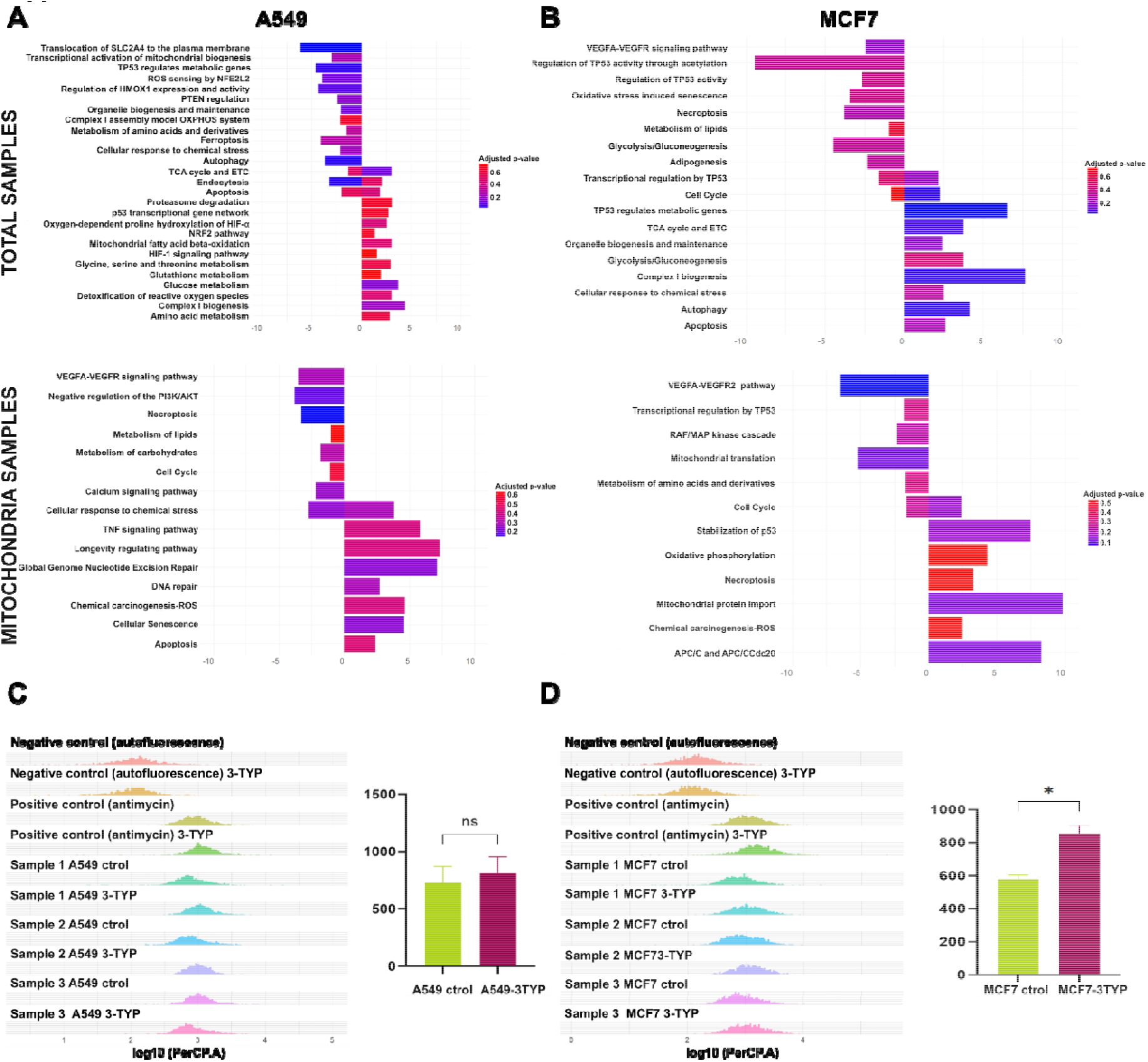
SIRT3 modulation uncovers distinct metabolic pathway enrichments in A549 and MCF7 cell lines. (A) Enriched biological pathways for total and mitochondrial proteomes in A549 cells, comparing untreated controls and 3-TYP–treated samples. (B) Equivalent analysis for MCF7 cells under the same conditions. Pathway enrichment was performed with GeneCodis using KEGG, Reactome, and WikiPathways annotations. (C) Flow cytometry histograms illustrating the data for the A549 cell line are displayed, comparing control and 3-TYP-treated samples. (D) Flow cytometry histograms for the MCF7 cell line are also shown, contrasting control conditions with those treated with 3-TYP.

In contrast, mitochondrial protein samples from the control group showed enrichment in pathways associated with tumor development. Among these, we identified the calcium signaling pathway, which is involved in the regulation of ion homeostasis; the VEGF signaling pathway, which is critical for angiogenesis and metastasis; and the necroptosis pathway (**Figure 3A**).

After treatment with 3-TYP, the total protein samples exhibited greater enrichment in TCA cycle and ETC pathways compared to the control group, along with increased enrichment in glucose metabolism and ROS detoxification pathways. Meanwhile, mitochondrial fractions from the treated group showed significant enrichment in DNA repair pathways, specifically nucleotide excision repair (GG-NER) (**Figure 3A**).

To validate the observed enrichment in ROS detoxification pathways under 3-TYP treatment, we conducted an assay to measure these species using MitoSOX, a reagent specifically designed to detect mitochondrial superoxide. However, we observed no significant differences between control and 3-TYP treated samples (**Figure 3C**).

Compared to the A549 cell line, the MCF7 cell line shows clear evidence of metabolic reprogramming and the tumor-suppressive role of SIRT3. Our proteomic analysis revealed that both control and 3-TYP treated total protein samples exhibited enrichment in pathways associated with glycolytic metabolism. However, after treatment with 3-TYP, a significant enrichment in oxidative metabolic pathways was observed, including the TCA cycle and the ETC (**Figure 3B**), while the OXPHOS pathway was particularly enriched in the mitochondrial fractions (**Figure 3B**).

Furthermore, the mitochondrial samples from the treated group showed significant enrichment in pathways related to the activation of the anaphase-promoting complex/cyclosome (APC/C and APC/CCdc20) (**Figure 3B**). APC/C, an E3 ubiquitin ligase, plays a crucial role in the degradation of key cell cycle proteins through its interaction with regulatory subunits Cdh1 and Cdc20. Notably, Cdc20 has been implicated in tumorigenesis and is recognized as an oncogene [23,24].

Although the MCF7 cell line exhibits reduced basal levels of SIRT3, its further inhibition with 3-TYP resulted in a significant increase in ROS, as evidenced by the MitoSOX assay (**Figure 3D**). This suggests that the additional loss of SIRT3 function is detrimental to this cell line.

### 3.4 Acetylation patterns observed in A549 and MCF7 cell lines suggests a role in ribosomal metabolism

In our analysis of acetylation stoichiometry, we established a threshold of 0.4% to define a peptide as hyperacetylated. When we evaluated the intrinsic differences in acetylation between the A549 and MCF7 cell lines, we identified 55 acetylated proteins in A549 and 56 in MCF7. Notably, only two proteins shared acetylation sites, as illustrated in **Figure 4A**.

**Figure 4.**
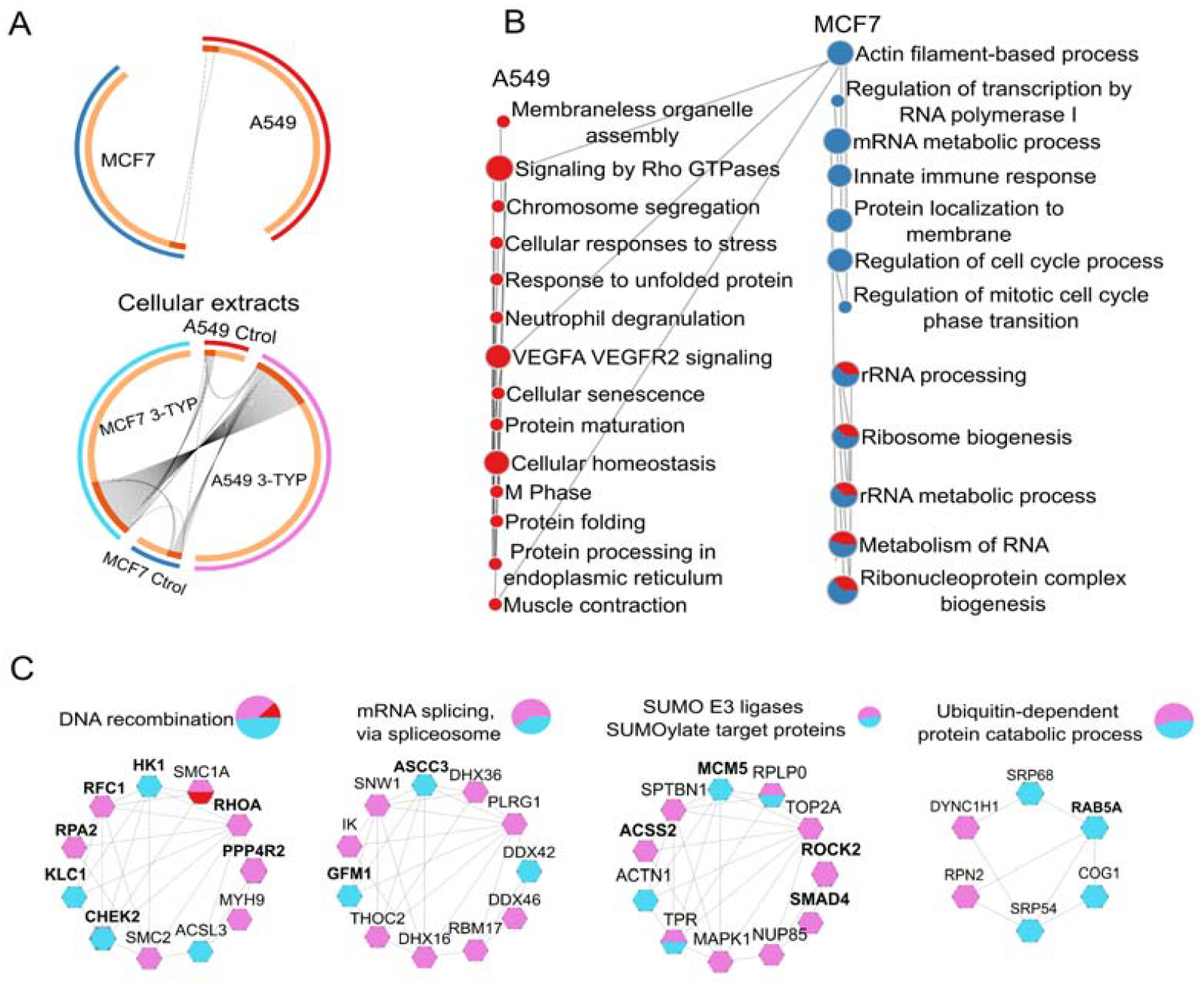
Acetylation patterns of total extracts from MCF7 and A549 cell lines. (A) The top panel compares hyperacetylated peptides in untreated A549 and MCF7 cell lines, with A549 control represented in red, while MCF7 is shown in blue. The bottom panel illustrates the comparison between treated and control samples: light blue indicates MCF7 treated with 3TYP, dark blue represents the MCF7 control, pink denotes A549 treated with 3TYP, and red indicates the A549 control. (B) This section highlights the enriched pathways associated with hyperacetylated proteins in the control samples of A549 (red) and MCF7 (blue). (C) The final section displays the enriched pathways for hyperacetylated proteins in treated versus control samples: light blue for treated MCF7, dark blue for MCF7 control, pink for treated A549, and red for A549 control.

To further investigate the functional implications of intrinsic acetylation in these cell lines, we performed a pathway enrichment analysis. We discovered that in both cell lines, acetylation was strongly associated with ribosomal RNA processing and ribosome biogenesis. However, we noted significant differences in the specific biological processes enriched in each cell type, reflecting the context-dependent role of SIRT3.

In the A549 lung cancer cells, where SIRT3 is proposed to function as an oncogene, hyperacetylated proteins were primarily involved in protein processing, cellular stress response, RHO GTPase signaling, and VEGFA signaling. This suggests a potential link between acetylation and tumor-promoting mechanisms, such as enhanced protein stability, increased cellular plasticity, and adaptation to stress. In contrast, in the MCF7 breast cancer cell line, acetylation was enriched in proteins related to cell cycle regulation, innate immune response modulation, and transcriptional regulation. This indicates that the role of acetylation in MCF7 cells may be more closely associated with maintaining genomic stability and controlling proliferative signals, consistent with the tumor-suppressive functions of SIRT3 (**Figure 4B**).

Treatment with 3-TYP revealed hyperacetylated proteins that could be key in cellular regulation. In A549 cells, hyperacetylation of the ACSS2 protein, a potential target of SIRT3, was identified, as well as in proteins involved in DNA replication and repair (**Figure 4C**).

In the MCF7 cell line, treatment with 3-TYP demonstrated notable hyperacetylation of the proteins KLC1 and RAB5A (Figure 4C), which are essential for maintaining cell morphology, cytoskeletal organization, and intracellular transport, factors crucial for preventing metastasis and preserving epithelial integrity. Additionally, hyperacetylation of the protein CHEK2, a key kinase in cell cycle regulation and DNA repair, was observed. Hyperacetylation of the mitochondrial protein ACSS3 was also noted, whose regulation is not fully understood, and whose activity appears to have opposing effects depending on the type of cancer [25–27].

### 3.5. SIRT3 regulates acetylation of metabolic proteins in A549 and MCF7 cells, affecting cancer progression

Mitochondrial acetylomes were analyzed independently for each cell line, comparing 3-TYP– treated samples with their respective controls. In A549 cells, SIRT3 inhibition resulted in 41 proteins meeting the predefined hyperacetylation threshold, whereas in MCF7 cells 224 proteins were classified as hyperacetylated (Figure 5A). This larger number in MCF7 suggests that acetylation homeostasis in this cell line may be more dependent on SIRT3 activity.

**Figure 5.**
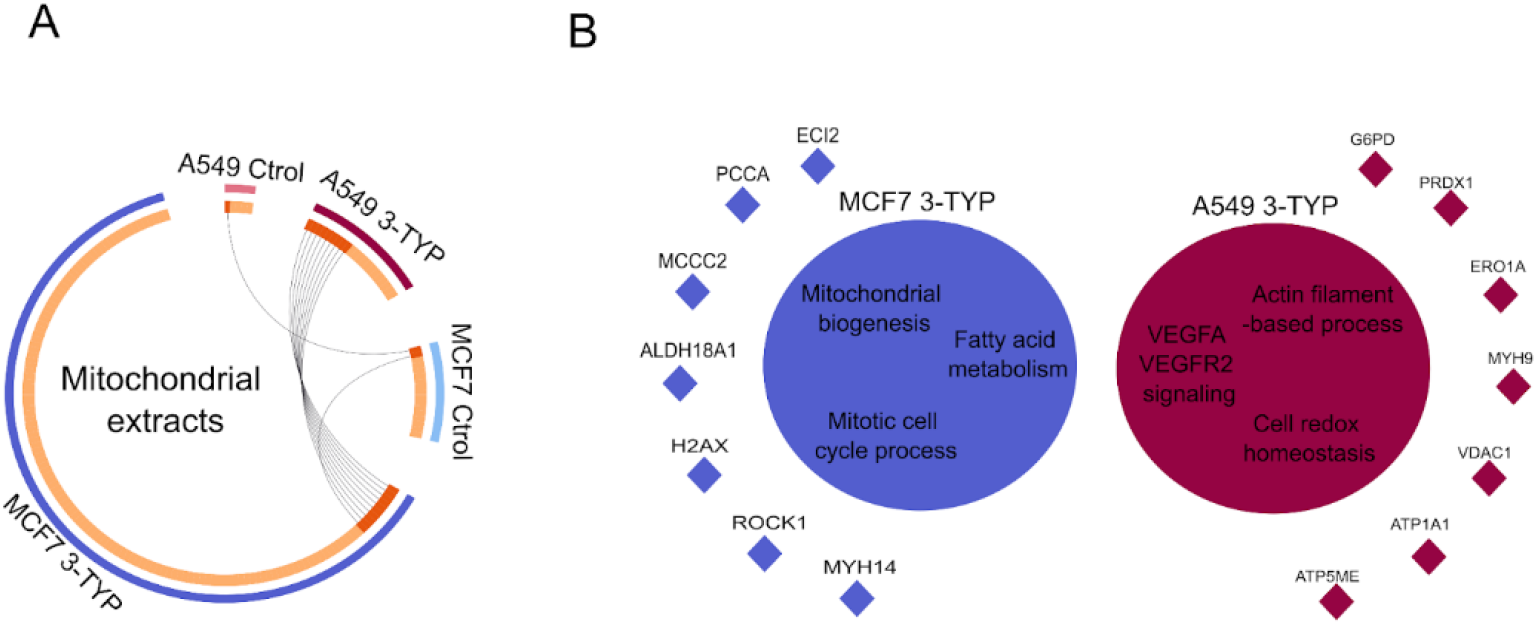
Acetylation profiles in mitochondrial extracts of MCF7 and A549 after SIRT3 inhibition. (A) Number and identity of mitochondrial proteins classified as hyperacetylated (acetylation stoichiometry ≥0.4%) after treatment with 3-TYP, shown separately for each cell line. Dark red: A549 treated; light red: A549 control; dark blue: MCF7 treated; light blue: MCF7 control. Proteins with hyperacetylated peptides detected in both cell lines are connected by black lines. Data represent within-cell-line comparisons (treated vs. control) and are not normalized between A549 and MCF7. (B) TPathway enrichment analysis of hyperacetylated proteins for each cell line, based on KEGG and Reactome annotations. Dark red: enriched pathways in 3-TYP–treated MCF7 vs. control; dark blue: enriched pathways in 3-TYP– treated A549 vs. control. Only pathways with adjusted P < 0.05 are shown.

In A549, hyperacetylated proteins were enriched in pathways related to VEGFA/VEGFR2 signaling, redox homeostasis, and maintenance of cell polarity, processes with established links to cancer biology (Figure 5B). Several of these proteins have not been extensively studied in the context of lysine acetylation, leaving their functional regulation by this modification unresolved. Among notable examples, VDAC1, an outer mitochondrial membrane channel regulating Ca^2+^ flux and linking glycolysis to oxidative phosphorylation via hexokinase binding, showed increased acetylation. Prior studies have identified VDAC1 as a SIRT3 deacetylation target. We also detected hyperacetylation in ATP5ME, a subunit of ATP synthase, and in ATP1A1, a Na^+^/K^+^-ATPase associated with tumor invasion and metastasis [28].

In MCF7, SIRT3 inhibition predominantly affected proteins involved in mitochondrial biogenesis, mitochondrial organization, and monocarboxylic acid catabolism. Notably, 3-methylcrotonyl-CoA carboxylase 2 (MCCC2), overexpressed in breast and colorectal cancers and linked to poor prognosis [29,30], was among the hyperacetylated proteins. Additional examples included PCCA and ECI2, whose acetylation status has been recently connected to altered tumor progression in different cancer models (Figure 5B). These findings point to candidate proteins whose acetylation changes may influence cancer-related phenotypes, although further functional validation will be required.

## 4. Discussion

The functional role of SIRT3 in NSCLC remains controversial. While some studies describe tumor-suppressive effects in adenocarcinoma cells, others report that SIRT3 overexpression can promote tumor progression and radioresistance [15,31].

A 2019 study by Pei-Hsuan and colleagues found a weak correlation between glucose consumption and growth rate in cancer cells, but a positive correlation with glutamine consumption, suggesting a preference for oxidative metabolism via OXPHOS [32]. Our proteomic results in the A549 cell line reflect a similar pattern, demonstrating enrichment of metabolic pathways in both total and mitochondrial protein control groups, with a clear inclination towards oxidative metabolism compared to the treated groups. Additionally, flow cytometry analysis of mitochondrial ROS generation revealed no significant changes after treatment with the specified inhibitor. Therefore, it seems that SIRT3 does not actively regulate the production of these species.

Acetylomic analysis revealed that treatment with the 3-TYP inhibitor promoted hyperacetylation of proteins involved in DNA replication and repair, in agreement with our proteomic findings. Among the proteins of interest, hyperacetylation of ACSS2, a potential target of SIRT3 [33], was particularly notable. Acetylation of ACSS2 leads to its inactivation, thereby preventing the synthesis of acetyl-CoA from acetate. This metabolic shift favors glycolysis over oxidative metabolism and may also reduce the mitochondrial pool of acetyl-CoA, consequently decreasing non-enzymatic acetylation.

Additionally, acetylation of ACSS2 at lysine 642 may inhibit its phosphorylation at serine 659 due to the close proximity of these two residues. Phosphorylation at this site is critical for the nuclear translocation of ACSS2, an event associated with poor cancer prognosis [34]. This interplay between acetylation and phosphorylation could represent a regulatory mechanism through which 3-TYP treatment modulates ACSS2 activity and subcellular localization, contributing to the metabolic and phenotypic changes observed in the studied cell line.

Regarding mitochondrial proteins, inhibition of SIRT3 in the A549 cell line led to an increase in hyperacetylated proteins enriched in pathways such as VEGFA/VEGFR2 signaling, cellular redox homeostasis, and maintenance of cell polarity. Among the hyperacetylated proteins of particular relevance were VDAC1, a regulator of mitochondrial communication and metabolism; ATP5ME, a component of the mitochondrial ATP synthase complex; and ATP1A1, a protein associated with tumor invasion. These findings suggest that SIRT3 may play a key role in regulating the acetylation of proteins involved in critical cellular processes related to cancer progression. However, further research is needed to elucidate the functional consequences of these acetylation events.

Due to the low expression of SIRT3 in the MCF7 cell line [17,18], treatment with the 3-TYP inhibitor did not result in significant changes in the enrichment of biological pathways. Nevertheless, it has been reported that total loss of SIRT3 function promotes a more aggressive phenotype in certain cancer types [35]. In line with this, our proteomic analysis revealed that, after treatment, pathways such as APC/C and APC/C–Cdc20 (linked to necroptosis) and VEGFA/VEGFR2 (associated with angiogenesis and vascular function) were enriched, all of which are closely related to poor cancer prognosis. These findings highlight the importance of SIRT3 function in this cell line.

SIRT3 inhibition also revealed a potential imbalance in oxidative metabolism pathways, evidenced by the increase in ROS. This ROS elevation may stabilize HIF-α, promoting the metabolic reprogramming of these cells and potentially impacting key cellular processes such as growth, proliferation, differentiation, and cell survival [36].

The acetylome of MCF7 cells treated with 3-TYP showed hyperacetylation of key metabolic proteins, among which we highlight hexokinase-1 (HK1), a crucial enzyme that channels glucose into glycolysis. GlcNAc-mediated inhibition of HK1 generates a feedback loop that balances glucose utilization across various metabolic pathways; therefore, its acetylation may enhance its activity, favoring greater glycolytic flux [37]. These findings underscore the pronounced metabolic shift in this cell line.

We also highlight the hyperacetylation of proteins KLC1 and RAB5A, which act synergistically to suppress breast cancer progression. KLC1, a subunit of the kinesin-1 motor complex, is mainly expressed in luminal/epithelial subtypes and inhibits epithelial-mesenchymal transition (EMT), invasion, metastasis, and the expression of stem cell markers, thus promoting a non-invasive tumor phenotype [38]. Meanwhile, RAB5A contributes to tumor suppression by facilitating the degradation of the epidermal growth factor receptor (EGFR), thereby attenuating PI3K/AKT/mTOR signaling, which is essential for cell survival, proliferation, and metastatic potential [39]. Additionally, we note the hyperacetylation of CHEK2, which has been associated with increased cancer risk due to its ability to stabilize the tumor suppressor p53 and interact with BRCA1 for homologous recombination-based DNA repair. Its acetylation may modulate its activity, affecting DNA damage response, genomic stability, and tumor progression. In mitochondrial samples treated with 3-TYP, we also found hyperacetylation in proteins that have been previously associated with cancer, such as MCCC2

A significant limitation of the methodology used to assess protein acetylation is the loss of information regarding low-abundance proteins. Given the importance of acetylation in global metabolism and cancer cell function, the lack of data on these proteins limits our understanding of the acetylome and its relationship to cancer. Therefore, it is necessary to explore and develop alternative methodologies to address these limitations and provide a more comprehensive view of the complex interactions between acetylation, metabolism, and cancer.

## 5. Conclusions

Within cell line proteomic and acetylomic analyses indicate that SIRT3 modulation produces distinct metabolic and signaling changes in A549 and MCF7 cells. In A549, higher basal SIRT3 levels were associated with oxidative metabolism. Inhibition with 3-TYP promoted the enrichment of pathways related to DNA repair, including nucleotide excision repair, which could indirectly improve metabolic efficiency by preserving mitochondrial and nuclear genome integrity. These findings suggest that in this context, SIRT3 inhibition might contribute to a more favorable metabolic state through the activation of repair mechanisms.

In MCF7, where SIRT3 expression is lower, inhibition induced more extensive hyperacetylation, enrichment of pathways associated with aggressive phenotypes, and increased ROS production. These changes, together with the acetylation of proteins involved in metabolic regulation, tumor suppression, and DNA damage response, indicate that further loss of SIRT3 function may be detrimental in this cellular context.

Overall, these results support the context dependent nature of SIRT3 function, shaped by the metabolic and molecular background of each cancer type. Potential therapeutic strategies targeting SIRT3 will require careful tailoring to the tumor’s intrinsic characteristics. Nevertheless, further functional studies are needed to support and expand these results.

## Acknowledgments

M.A-R, was a master degree student in the Programa de Maestría en Ciencias Bioquímicas, Universidad Nacional Autónoma de México (UNAM). She has been awarded a fellowship from the Consejo Nacional de Ciencias Humanidades, y Tecnología (CONAHCyT) with the reference number 1145601 We would like to express our gratitude to Erika Isabel Melchy Perez and Angel Gabriel Martínez-Batallar for their technical support.

## Data Availability Statement

The mass spectrometry proteomics data are available to the ProteomeXchange Consortium via the PRIDE partner repository with the data set identifier PXD063181. Any additional requests can be directed to the corresponding author.

## References

1. Memon AA, Vats S, Sundquist J, Li Y, Sundquist K. Mitochondrial DNA Copy Number: Linking Diabetes and Cancer. Antioxid Redox Signal. 2022;37: 1168–1190.

2. Gil J, Ramírez-Torres A, Encarnación-Guevara S. Lysine acetylation and cancer: A proteomics perspective. J Proteomics. 2017;150: 297–309.

3. Narita T, Weinert BT, Choudhary C. Functions and mechanisms of non-histone protein acetylation. Nat Rev Mol Cell Biol. 2019;20: 156–174.

4. Wagner GR, Payne RM. Widespread and enzyme-independent Nε-acetylation and Nε-succinylation of proteins in the chemical conditions of the mitochondrial matrix. J Biol Chem. 2013;288: 29036–29045.

5. Lombard DB, Alt FW, Cheng H-L, Bunkenborg J, Streeper RS, Mostoslavsky R, et al. Mammalian Sir2 homolog SIRT3 regulates global mitochondrial lysine acetylation. Mol Cell Biol. 2007;27: 8807–8814.

6. Sirtuin 3 (SIRT3) Protein Regulates Long-chain Acyl-CoA Dehydrogenase by Deacetylating Conserved Lysines Near the Active Site. Journal of Biological Chemistry. 2013;288: 33837–33847.

7. Zhang Y, Bharathi SS, Rardin MJ, Uppala R, Verdin E, Gibson BW, et al. SIRT3 and SIRT5 Regulate the Enzyme Activity and Cardiolipin Binding of Very Long-Chain Acyl-CoA Dehydrogenase. PLOS ONE. 2015;10: e0122297.

8. Tyr Phosphorylation of PDP1 Toggles Recruitment between ACAT1 and SIRT3 to Regulate the Pyruvate Dehydrogenase Complex. Molecular Cell. 2014;53: 534–548.

9. SIRT3 Protein Deacetylates Isocitrate Dehydrogenase 2 (IDH2) and Regulates Mitochondrial Redox Status. Journal of Biological Chemistry. 2012;287: 14078–14086.

10. A small molecule activator of SIRT3 promotes deacetylation and activation of manganese superoxide dismutase. Free Radical Biology and Medicine. 2017;112: 287–297.

11. Advances in characterization of SIRT3 deacetylation targets in mitochondrial function. Biochimie. 2020;179: 1–13.

12. Baeza J, Smallegan MJ, Denu JM. Mechanisms and Dynamics of Protein Acetylation in Mitochondria. Trends Biochem Sci. 2016;41: 231–244.

13. Chen Y, Fu LL, Wen X, Wang XY, Liu J, Cheng Y, et al. Sirtuin-3 (SIRT3), a therapeutic target with oncogenic and tumor-suppressive function in cancer. Cell Death & Disease. 2014;5: e1047–e1047.

14. Bray F, Laversanne M, Sung H, Ferlay J, Siegel RL, Soerjomataram I, et al. Global cancer statistics 2022: GLOBOCAN estimates of incidence and mortality worldwide for 36 cancers in 185 countries. CA Cancer J Clin. 2024;74: 229–263.

15. Cao K, Chen Y, Zhao S, Huang Y, Liu T, Liu H, et al. Sirt3 Promoted DNA Damage Repair and Radioresistance Through ATM-Chk2 in Non-small Cell Lung Cancer Cells. J Cancer. 2021;12: 5464–5472.

16. Xiong Y, Wang M, Zhao J, Wang L, Li X, Zhang Z, et al. SIRT3 is correlated with the malignancy of non-small cell lung cancer. Int J Oncol. 2017;50: 903–910.

17. Park JS, Lee S, Jeong AL, Han S, Ka HI, Lim J-S, et al. Hypoxia-induced IL-32β increases glycolysis in breast cancer cells. Cancer Lett. 2015;356: 800–808.

18. Schwab LP, Peacock DL, Majumdar D, Ingels JF, Jensen LC, Smith KD, et al. Hypoxia-inducible factor 1α promotes primary tumor growth and tumor-initiating cell activity in breast cancer. Breast Cancer Res. 2012;14: R6.

19. Finley LWS, Carracedo A, Lee J, Souza A, Egia A, Zhang J, et al. SIRT3 opposes reprogramming of cancer cell metabolism through HIF1α destabilization. Cancer Cell. 2011;19: 416–428.

20. Garcia-Moreno A, López-Domínguez R, Villatoro-García JA, Ramirez-Mena A, Aparicio-Puerta E, Hackenberg M, et al. Functional Enrichment Analysis of Regulatory Elements. Biomedicines. 2022;10. doi:10.3390/biomedicines10030590

21. Zhang J, Xiang H, Liu J, Chen Y, He R-R, Liu B. Mitochondrial Sirtuin 3: New emerging biological function and therapeutic target. Theranostics. 2020;10: 8315–8342.

22. Pinterić M, Podgorski II, Hadžija MP, Filić V, Paradžik M, Proust BLJ, et al. Sirt3 Exerts Its Tumor-Suppressive Role by Increasing p53 and Attenuating Response to Estrogen in MCF-7 Cells. Antioxidants (Basel). 2020;9. doi:10.3390/antiox9040294

23. Schrock MS, Stromberg BR, Scarberry L, Summers MK. APC/C ubiquitin ligase: Functions and mechanisms in tumorigenesis. Semin Cancer Biol. 2020;67: 80–91.

24. Greil C, Engelhardt M, Wäsch R. The Role of the APC/C and Its Coactivators Cdh1 and Cdc20 in Cancer Development and Therapy. Front Genet. 2022;13: 941565.

25. Chang W-C, Cheng W-C, Cheng B-H, Chen L, Ju L-J, Ou Y-J, et al. Mitochondrial Acetyl-CoA Synthetase 3 is Biosignature of Gastric Cancer Progression. Cancer Med. 2018;7: 1240–1252.

26. Sui L, Sanders A, Jiang WG, Ye L. Deregulated molecules and pathways in the predisposition and dissemination of breast cancer cells to bone. Comput Struct Biotechnol J. 2022;20: 2745–2758.

27. Zhou L, Song Z, Hu J, Liu L, Hou Y, Zhang X, et al. ACSS3 represses prostate cancer progression through downregulating lipid droplet-associated protein PLIN3. Theranostics. 2021;11: 841–860.

28. Chen Y-I, Chang C-C, Hsu M-F, Jeng Y-M, Tien Y-W, Chang M-C, et al. Homophilic ATP1A1 binding induces activin A secretion to promote EMT of tumor cells and myofibroblast activation. Nat Commun. 2022;13: 2945.

29. Dai W, Feng H, Lee D. MCCC2 overexpression predicts poorer prognosis and promotes cell proliferation in colorectal cancer. Exp Mol Pathol. 2020;115: 104428.

30. Liu Y, Yuan Z, Song C. Methylcrotonoyl-CoA carboxylase 2 overexpression predicts an unfavorable prognosis and promotes cell proliferation in breast cancer. Biomark Med. 2019;13: 427–436.

31. Xiong Y, Wang L, Wang S, Wang M, Zhao J, Zhang Z, et al. SIRT3 deacetylates and promotes degradation of P53 in PTEN-defective non-small cell lung cancer. J Cancer Res Clin Oncol. 2018;144: 189–198.

32. Chen P-H, Cai L, Huffman K, Yang C, Kim J, Faubert B, et al. Metabolic Diversity in Human Non-Small Cell Lung Cancer Cells. Mol Cell. 2019;76: 838–851.e5.

33. Hallows WC, Lee S, Denu JM. Sirtuins deacetylate and activate mammalian acetyl-CoA synthetases. Proc Natl Acad Sci U S A. 2006;103: 10230–10235.

34. Yang X, Shao F, Shi S, Feng X, Wang W, Wang Y, et al. Prognostic Impact of Metabolism Reprogramming Markers Acetyl-CoA Synthetase 2 Phosphorylation and Ketohexokinase-A Expression in Non-Small-Cell Lung Carcinoma. Front Oncol. 2019;9: 1123.

35. Giralt A, Villarroya F. SIRT3, a pivotal actor in mitochondrial functions: metabolism, cell death and aging. Biochem J. 2012;444: 1–10.

36. Storz P. Reactive oxygen species in tumor progression. Front Biosci. 2005;10: 1881–1896.

37. Wolf AJ, Reyes CN, Liang W, Becker C, Shimada K, Wheeler ML, et al. Hexokinase Is an Innate Immune Receptor for the Detection of Bacterial Peptidoglycan. Cell. 2016;166: 624–636.

38. A role for kinesin-1 subunits KIF5B/KLC1 in regulating epithelial mesenchymal plasticity in breast tumorigenesis. EBioMedicine. 2019;45: 92–107.

39. NAA20 recruits Rin2 and promotes triple-negative breast cancer progression by regulating Rab5A-mediated activation of EGFR signaling. Cellular Signalling. 2023;112: 110922.

